# Innate immune signaling drives late cardiac toxicity following DNA damaging cancer therapies

**DOI:** 10.1101/2021.09.26.461865

**Authors:** Achraf Shamseddine, Suchit H. Patel, Valery Chavez, Mutayyaba Adnan, Melody Di Bona, Jun Li, Chau T. Dang, Lakshmi V. Ramanathan, Kevin C. Oeffinger, Jennifer E. Liu, Richard M. Steingart, Alessandra Piersigilli, Nicholas D. Socci, Angel T. Chan, Anthony F. Yu, Samuel F. Bakhoum, Adam M. Schmitt

**Affiliations:** Division of Translational Oncology, Department of Radiation Oncology, Memorial Sloan Kettering Cancer Center; New York, NY, USA; Department of Radiation Oncology, Mary Bird Perkins Cancer Center; Baton Rouge, LA, USA; Human Oncology and Pathogenesis Program, Memorial Sloan Kettering Cancer Center; New York, NY, USA; Breast Medicine Service, D Department of Medicine, Memorial Sloan Kettering Cancer Center; New York, NY, USA; Clinical Chemistry Service, Department of Laboratory Medicine, Memorial Sloan Kettering Cancer Center; New York, NY, USA; Department of Medicine, Duke University School of Medicine, Durham, NC, USA; Cardiology Service, Department of Medicine, Memorial Sloan Kettering Cancer Center; New York, NY, USA; Laboratory of Comparative Pathology, Rockefeller University, Weill Cornell Medicine and Memorial Sloan-Kettering Cancer Center, New York, New York, USA; Marie-Josee & Henry R. Kravis Center for Molecular Oncology, Memorial Sloan Kettering Cancer Center, New York, NY, USA

## Abstract

Late cardiac toxicity is a potentially lethal complication of cancer therapy, yet the pathogenic mechanism remains largely unknown, and few treatment options exist. Here we report DNA damaging agents such as radiation and anthracycline chemotherapies induce delayed cardiac inflammation following therapy due to activation of cGAS and STING-dependent type I interferon signaling. Genetic ablation of cGAS-STING-signaling in mice inhibits DNA damage induced cardiac inflammation, rescues late cardiac functional decline, and prevents death from cardiac events. Treatment with a STING antagonist suppresses cardiac interferon signaling following DNA damaging therapies and effectively mitigates cardiotoxicity. These results identify a therapeutically targetable, pathogenic mechanism for one of the most vexing treatment-related toxicities in cancer survivors.

## Main Text

Radiation therapy (RT) and anthracycline chemotherapeutics such as doxorubicin are central to the curative treatment regimens of many cancers. These treatments elicit cytotoxic effects in cancer cells through creation of DNA double stranded breaks and are often used individually or in combination to treat cancer of the breast cancer, thorax, and Hodgkin’s and Non-Hodgkin’s lymphomas ^1-3^. As cancer long-term survival continues to increase, late normal tissue toxicities, especially late cardiac toxicity, are a major source of long-term morbidity and mortality in cancer survivors ^4,5^. As such, there is a growing need to identify mechanisms of late normal tissue toxicity in order to develop therapeutic strategies to mitigate these late effects. RT-induced cardiac toxicity doubles with every Gray of mean heart dose following treatment for breast cancer ^6^ and its lifetime occurrence was found to be at 8% in long term survivors of Hodgkin’s lymphoma, fivefold higher than the general population ^7^. Similarly, the risk of late doxorubicin-induced cardiotoxicity exceeds 35% with cumulative doses over 600 mg/m^2 8^.

While the acute effects of DNA double stranded breaks from RT and anthracyclines are well known, the mechanisms by which DNA damage leads to normal tissue remodeling and late tissue toxicity years after cancer treatment are poorly understood ^9^. The generation of reactive oxygen species and topoisomerase-2β-mediated mitochondrial dysfunction and apoptosis of cardiomyocytes following treatment have been implicated in pathogenesis of cardiac toxicity following RT and doxorubicin ^10,11^. Concurrent administration of dexrazoxane, an iron chelator that reduces superoxide radical generation and inhibits topoisomerase-2β, is an FDA approved therapy for prevention of doxorubicin associated cardiac toxicity ^12^. However, the intervening mechanisms connecting acute tissue stress to delayed and chronic tissue damage are poorly understood. Experimental data have shown that DNA damage induces sterile inflammation in tumors and in tissue ^13^. However, studies of inflammatory signaling following DNA damaging therapies have predominantly focused on transforming growth factor β (TGF-β) and downstream pro-inflammatory cytokines, and have yet to translate into effective strategies to mitigate late normal tissue toxicity.

Since late normal tissue toxicity likely stems from a combination of cell intrinsic and microenvironmental responses to cardiac DNA damage, we developed an animal model of DNA damage-induced cardiac toxicity to examine the tissue-specific temporal response to cardiac DNA damage. Mice were treated with a single dose of cardiac RT using an image guided radiotherapy platform to isolate high dose exposure to the heart and minimize dose to other organs. Mice were euthanized at timepoints from 6 hours to 28 days following treatment with a single dose of 12Gy RT, at which time hearts tissues were minced and dissociated to single cells suspensions, and cardiomyocytes, endothelia, and fibroblasts were isolated and RNA extracted for gene expression analyses by RNA-seq (**Figure 1a**). Comparison of tissue specific gene expression across specimens confirmed successful enrichment for each tissue population (**Figure 1b**). The acute response to cardiac DNA damage at 6 hours was similar across all three cell populations, driven by expression of DNA damage response genes, p53-dependent genes, and cell cycle regulators, all of which subsequently waned by 28 days following DNA damage (**Figure 1c, Supplementary Table 1**). In contrast, gene ontology enrichment analysis revealed substantial enrichment of genes associated with interferon responses and anti-viral signaling, such as *Irf7*, amongst genes induced 28 days after RT, a response that was largely restricted to cardiac fibroblasts (**Figures 1c-e**). We next examined whether DNA damage-induced interferon signaling in hearts also occurred in response to DNA damage by anthracyclines. Cardiac endothelial cells and fibroblasts were isolated from mice two weeks after completion of treatment with 5 weekly doses of doxorubicin (**Figure 1f**), a regimen previously shown to induce cardiac toxicity ^10^. Similar to mice after cardiac RT, cardiac fibroblasts, but not cardiac endothelial cells, responded to DNA damage with delayed activation of *Irf7* (**Figure 1g**). Therefore, delayed activation of interferon stimulated genes specifically in cardiac fibroblasts is a common response to heart exposure to DNA damaging agents.

**Figure 1.**
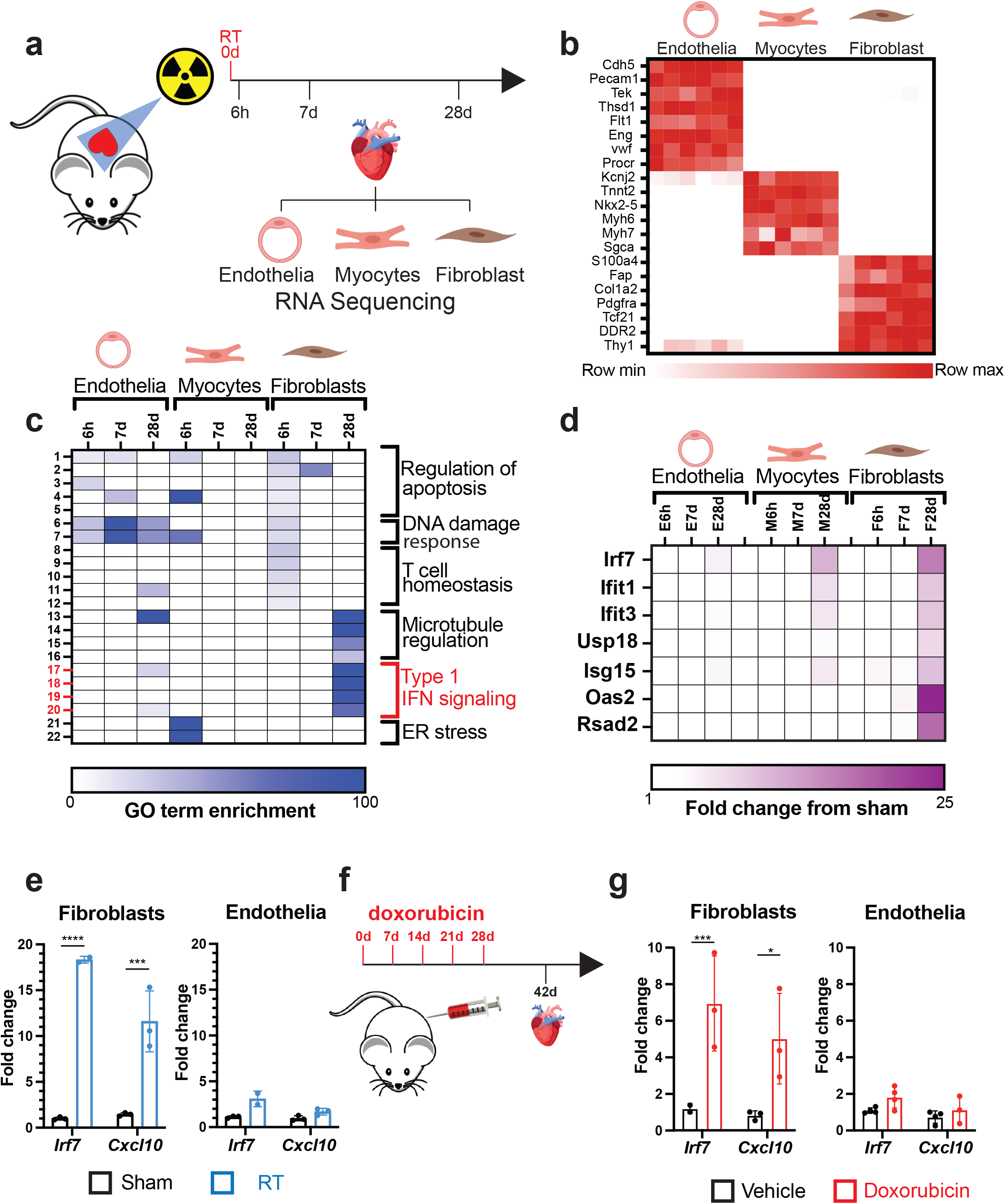
Delayed activation of type I interferon signaling after cardiac DNA damage. **a**, Schematic representation of experimental model of radiation induced cardiac toxicity. **b**, Heatmap of gene expression of cell-specific markers in endothelia, myocytes and fibroblasts quantified by RNA-seq. **c**, Heatmap of GO term fold enrichment in endothelia, myocytes and fibroblasts at 6 hours, 7 days and 28 days after cardiac RT. **d**, Expression heatmap of type I-interferon-related genes at 6 hours, 7 days and 28 days post-RT in endothelial cells, myocytes and fibroblasts. **e**, Gene expression of *Irf7* and *Cxcl10* measured by qRT-PCR in endothelial cells and fibroblasts for 28 days after 12Gy cardiac RT or sham treatment. **f**, Schematic representation of mouse model of doxorubicin induced cardiac toxicity. **g**, qRT-PCR of endothelial cells and fibroblasts for *Irf7* and *Cxcl10* 14 days after completing 5 weekly doses of doxorubicin or vehicle only. *p<0.05, **p<0.01, ***p<0.001, ****p<0.0001.

Given the strong activation of a pro-inflammatory, type I interferon response following RT and doxorubicin, we sought to determine the mechanism of this activation. Multiple endogenous molecules, collectively referred to as damage-associated molecular patterns (DAMPs) have been shown to drive a robust interferon activation ^14^. Given the DNA damaging action of both radiation and doxorubicin, we reasoned that this type I interferon response may be driven by intracellular recognition of damaged nucleic acid through intracellular pattern recognition receptors (PRRs) ^15,16^. Mice deficient in cytosolic double stranded RNA (dsRNA) signaling (*Mavs*^*-/-*^) still exhibited a strong *Irf7* induction following RT, suggesting that detection of cytosolic dsRNA is not the driver of response (**Figure 2a**). However, ablation of cytosolic dsDNA recognition pathway in *Cgas*^*-/-*^ and *Sting*^*gt/gt*^ mice completely prevented *Irf7 and Cxcl10* induction following RT (**Figures 2a-b**). Similarly, absence of *Irf7 and Cxcl10* induction was noted in *Sting*^*gt/gt*^ mice following doxorubicin administration (**Figures 2c**). Pathogenic cytosolic DNA is detected by cGAS which signals to STING via production of the second messenger cyclic GMP-AMP ^17^. Prior reports have revealed that recruitment of cGAS to cytosolic DNA generate cGAS foci ^18,19^. Since cGAS-STING-dependent interferon-stimulated genes (ISG) induction following DNA damage was restricted largely to cardiac fibroblasts, we next examined cardiac tissue for cytosolic cGAS foci following RT. Using immunofluorescence imaging of heart tissue 28 days following RT, both nuclear and cytosolic cGAS foci were increased in vimentin-positive cells, a marker of cardiac fibroblasts, but not in vimentin-negative cells (**Figures 2d-g, Supplementary Figures 1a-b**).

**Figure 2.**
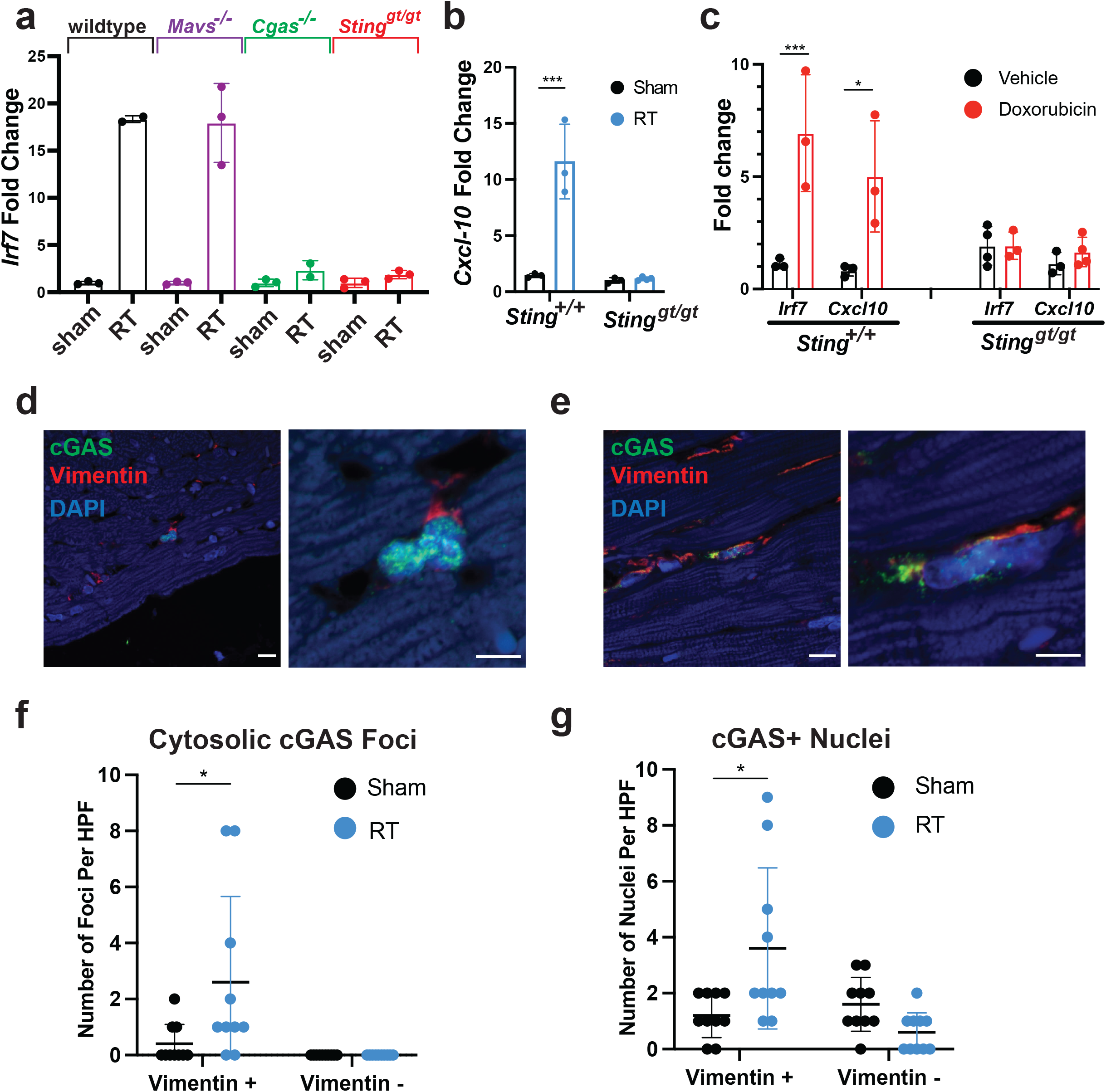
Type I interferon signaling after DNA damage requires cGAS and STING. **a**, *Irf7* expression measured by qRT-PCR in fibroblasts of mice genetically deficient in different components of PRR signaling, 28 days after cardiac RT. **b**, *Cxcl10* expression measured by qRT-PCR in fibroblasts of *Sting*^*+/+*^ vs *Sting*^*gt/gt*^ 28 days after cardiac RT. **c**, *Irf7* and *Cxcl10* expression measured by qRT-PCR in fibroblasts of *Sting*^*+/+*^ vs *Sting*^*gt/gt*^ 14 days after completing 5 weekly doses of doxorubicin. **d**, Immunofluorescence for cGAS (green) and Vimentin (red) in heart sections of *Sting*^*+/+*^ mice 28 days after sham treatment. **e**, Immunofluorescence for cGAS (green) and Vimentin (red) in heart sections *Sting*^*+/+*^ mice 28 days after 20Gy cardiac RT. **f**, Quantification of the number of cells with cytosolic cGAS-positive foci in sham-treated and RT-treated *Sting*^*+/+*^ mice. Each measurement represents number of cells in one high powered field of view. **g**, Quantification of the number of cells with cGAS positive nuclei in sham-treated and RT-treated *Sting*^*+/+*^ mice. Each measurement represents number of cells in one high powered field of view. *p<0.05, **p<0.01, ***p<0.001, ****p<0.0001.

Expression of ISGs raised the possibility that interferon signaling could remodel the inflammatory infiltrate in hearts following DNA damaging cancer therapy. We performed single cell RNA-seq on CD45+ leukocytes isolated from the mouse myocardium to determine whether cardiac DNA damage and ensuing cGAS-STING signaling cooperate to remodel the leukocyte repertoire in the heart. At 28 days after cardiac RT, when cGAS foci and ISG induction is apparent in cardiac fibroblasts, IFN activated monocytes and macrophages are recruited to the myocardium of wildtype mice, but this response is attenuated in the hearts of *Sting*^*gt/gt*^ mice (**Figure 3a-b, Supplementary Figure 2, Supplementary Table 2**). An identical cell population is recruited to the hearts of mice following myocardial infarction and has been associated with pathologic cardiac remodeling leading to loss of cardiac function and worsened survival ^20,21^. Therefore, cGAS-STING-dependent type I IFN signaling induced by cardiac DNA damage drives inflammation in the cardiac microenvironment which is associated with cardiac pathologic processes and death from cardiac events.

**Figure 3.**
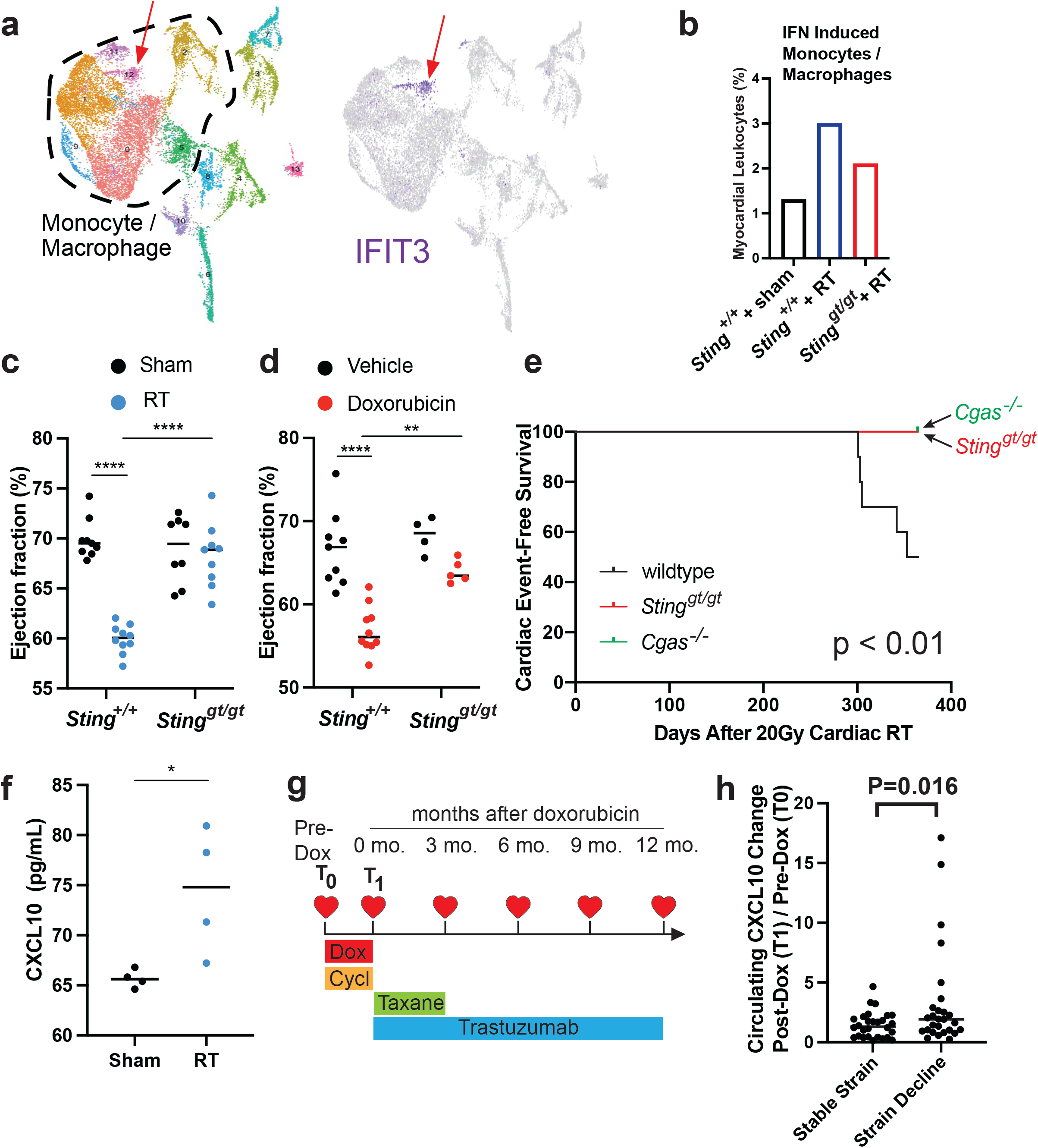
cGAS-STING signaling drives cardiac toxicity after DNA damaging cancer therapy. **a**, Left: UMAP defined clusters of cells from scRNA-seq of CD45-positive leukocytes isolated from the myocardium of mice 28 days after cardiac RT. Right: UMAP plot overlaid with the expression of the type I interferon stimulated gene *Ifit3*. **b**, Cluster 12 (IFN induced macrophages and monocytes) as a percentage of the total cell population in *Sting*^*+/+*^ after 28 days after sham treatment, *Sting*^*+/+*^ mice 28 days after cardiac RT, and *Sting*^*gt/gt*^ mice 28 days after cardiac RT. **c**, Ejection fraction measured by 2D-echocardiography in *Sting*^*+/+*^ vs *Sting*^*gt/gt*^ mice 3 months after cardiac RT. **d**, Ejection fraction measured by 2D-echocardiography *Sting*^*+/+*^ vs *Sting*^*gt/gt*^ mice 14 days after completing 5 weekly doses of doxorubicin. **e**, Kaplan Meier plot of survival of wildtype (n=10), *Cgas*^*-/-*^ (n=12) and *Sting*^*gt/gt*^ (n=12) mice after cardiac RT, day 0 is denoted as day of cardiac RT, tick marks indicate that 12 of 12 *Cgas*^*-/-*^ and 12 of 12 *Sting*^*gt/gt*^ remain alive 365 days after cardiac RT. **f**, Circulating CXCL10 chemokine levels in mice 28 days after cardiac RT or sham treatment, as measured by ELISA. **g**, Study schema for MSKCC IRB 14-099, a clinical trial to identify biomarkers of cardiac toxicity in HER2+ breast cancer patients receiving doxorubicin based polychemotherapy and trastuzumab. Timepoints of echocardiograms and blood biospecimens obtained on the trial in relation to the treatment schedule. T_0_ denotes pre-treatment timepoint and T_1_ denotes a post-doxorubicin, pre-trastuzumab timepoint. **h**, The change in circulating CXCL10 levels from T_0_ to T_1_ levels in patients who developed >15% decline in GLS, a early imaging biomarker of cardiac toxicity, compared to those who did not. *p<0.05, **p<0.01, ***p<0.001, ****p<0.0001.

We next evaluated whether this signaling pathway also contributed to physiologic cardiac parameters following DNA damaging cancer therapies. Cardiac functional decline as measured by echocardiogram is an indication of cardiac toxicity from cancer therapy. While wildtype mice treated with cardiac RT to 20Gy had no discernable change in cardiac function one month following RT, by 3 months irradiated animals began to show signs of cardiac dysfunction with declines in ejection fraction (EF) (**Figure 3c**) and diminished fractional shortening (FS) in comparison to sham irradiated wildtype mice of the same age (**Supplementary Figure 3a**). In contrast, identical cardiac irradiation of *Sting*^*gt/gt*^ mice demonstrated no sign of cardiac functional decline compared to age matched sham irradiated mice (**Figure 3c, Supplementary Figure 3a**). Doxorubicin treated wildtype mice experience reduced EF, decrease FS, and increased end diastolic volume compared to sham treated mice 2 weeks after completing a five-dose regimen of doxorubicin, while loss of cGAS-STING signaling in *Sting*^*gt/gt*^ mice rescued cardiac function, similar to results in mice treated with cardiac RT (**Figure 3d, Supplementary Figure 3b**). Furthermore, cardiac functional decline at 3 months following cardiac RT was predictive of survival by one year after irradiation. Death from cardiac events in wildtype mice (n=10) occurred beginning 300 days after RT with 50% incidence of death from cardiac events by 1 year (**Figure 3e**). Cardiac events were associated uniformly with signs of severe respiratory distress prior to death and pleural effusions at the time of necropsy. Consistent with cardiac toxicity, histopathology of the heart revealed cardiomyocyte degeneration, fibrosis, and luminal thrombi. In marked contrast, 100% of *Cgas*^*-/-*^ (n=12) and *Sting*^*gt/gt*^ (n=12) mice survived at 365 days following cardiac RT with no signs of cardiac toxicity (**Figure 3e**). Therefore, cGAS-STING signaling initiated in the weeks following cardiac DNA damage drives functional cardiac decline and death from late cardiac toxicity months after treatment.

Since cardiac DNA damage induces the expression of *Cxcl10* in fibroblasts in a cGAS-STING-dependent manner and circulating levels of CXCL10 have previously been associated with cardiovascular disease ^22^, we next examined whether cardiac DNA damage increases the circulating levels of CXCL10 in mice. CXCL10 is significantly increased in the blood of mice receiving cardiac RT 28 days after treatment relative to sham treated mice of the same age (**Figure 3f**), corresponding to the time at which *Cxcl10* is induced in a cGAS-STING-dependent manner after RT (**Figure 2a**). Since circulating CXCL10 is induced by cardiac DNA damage, we next queried whether CXCL10 dynamics following cardiac DNA damage was associated with clinical correlates of cardiac toxicity. In cancer patients treated with cardiotoxic therapies such as RT or anthracyclines, a decline in global longitudinal strain (GLS) is considered an early imaging biomarker of cardiac toxicity that precedes overt cardiac functional decline. Therefore, we next examined the association of CXCL10 with changes in GLS in cancer patients receiving cardiotoxic therapy. MSKCC IRB 14-099 enrolled 80 HER2+ breast cancer patients receiving standard of care anthracyclines based polychemotherapy followed by trastuzumab to monitor for cardiac toxicity using serial echocardiograms and to obtain blood specimens for identification of predictive biomarkers of cardiotoxicity. In this trial, 45% of patients developed a GLS decline >15% within one year of completing doxorubicin. Using these biospecimens, we quantified the circulating CXCL10 levels at pre-treatment (T_0_) and post-anthracycline, pre-trastuzumab (T_1_) and examined their relationship to GLS decline (**Figure 3g**). Specimens were available at T_0_ and T_1_ for 56 of these patients, of whom 27 (48%) had >15% decline in GLS within one year of completing doxorubicin while 29 (52%) did not. The change in circulating CXCL10 levels from (T_0_) to (T_1_) was significantly higher in patients who subsequently experience a decline in GLS compared to those who did not (**Figure 3h**). Since these data points precede exposure to paclitaxel or trastuzumab, they are thus solely a metric of a patient’s inflammatory response to a 2-month regimen of DNA damaging chemotherapy including doxorubicin and cyclophosphamide and are consistent with our results in mouse models that activation of type I IFN signaling after cardiac DNA damaging therapy drives late cardiac toxicity.

The cGAS-STING pathway is the subject of intensive pre-clinical drug development in oncology and autoimmune diseases and several small molecule antagonists have recently been developed. We next wanted to test whether pharmacologic inhibition of the cGAS-STING pathway is an effective therapeutic approach for the prevention of cardiac toxicity following DNA damaging cancer therapy. The recent characterization of small molecule covalent inhibitors of STING, such as H-151 and C-176, offered an attractive tool ^23^. Since we observed that DNA damage-induced, cGAS-STING-dependent type I interferon signaling was detectable by 28 days following RT, we began daily treatments with H-151 21 days after RT and continued until the end point of the study (**Figure 4a**). H-151 treatments effectively attenuated *Irf7* and *Cxcl-10* induction 28 days after RT in cardiac fibroblasts of wildtype animals, reducing levels similar to those in *Sting*^*gt/gt*^ mice (**Figure 4b**). C-176 treatments on the other hand had a less robust effect on *Irf7* and *Cxcl-10* following RT (**Supplementary Figures 4a-b**) Having confirmed the activity of H-151 to inhibit cGAS-STING signaling *in vivo* in our model of cardiac toxicity, we next examined whether H-151 was capable of preventing cardiac functional decline following cardiac DNA damage. Echocardiography of wildtype mice treated with H-151 or vehicle alone showed that H-151 prevented a decline in EF and FS in wildtype animals following RT (**Figure 4c, Supplementary Figure 4c**). Similarly, in doxorubicin treated mice, H-151 treatments prevented ejection fraction decline, as well as ventricular dilation manifested through increase in left ventricular end diastolic volume (**Figures 4d-e, Supplementary Figure 4d**). Therefore, cGAS-STING pathway antagonists are promising therapies for prevention of cardiac toxicity following cancer therapies that cause cardiac DNA damage.

**Figure 4.**
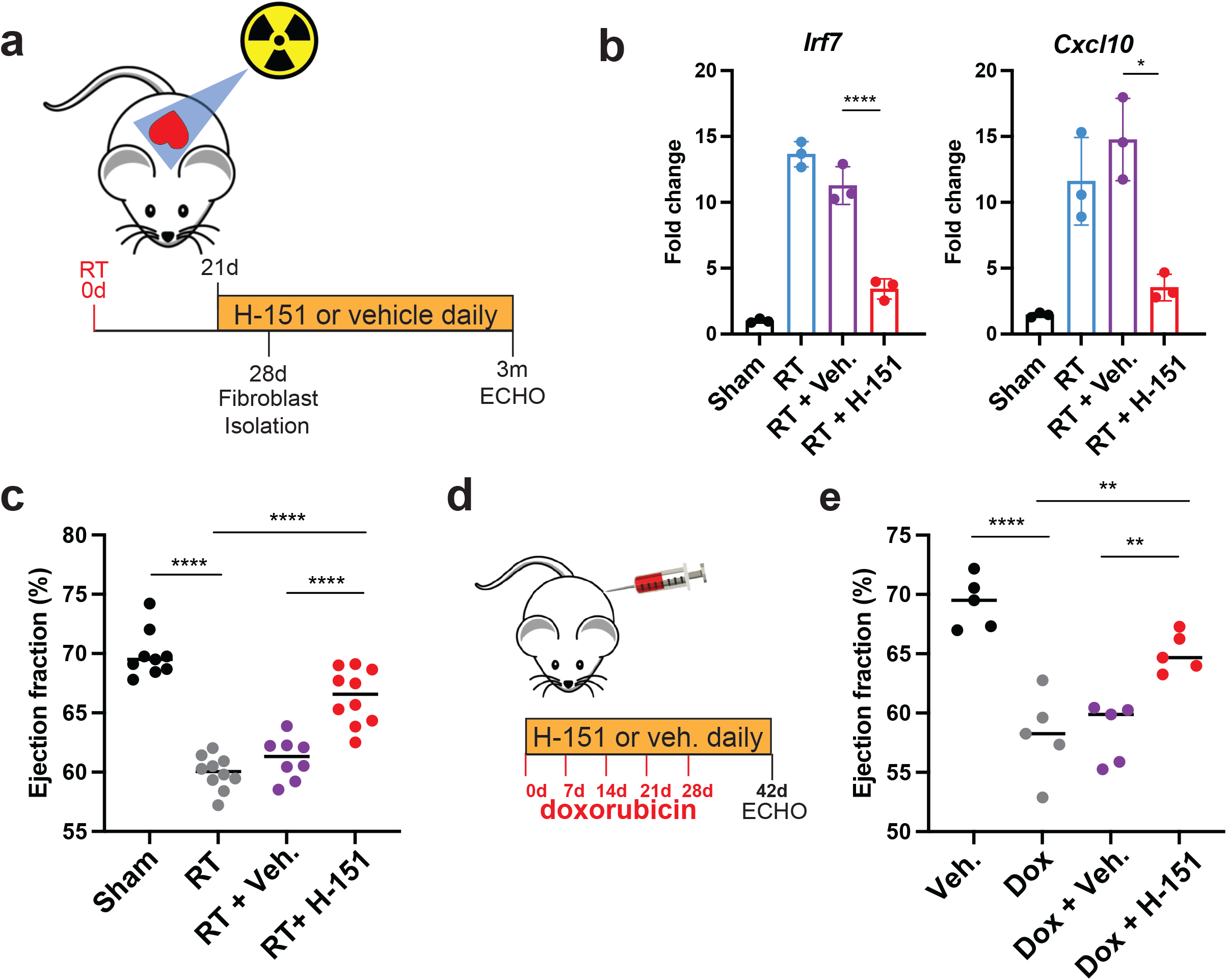
A covalent STING inhibitor effectively prevents cardiac toxicity from RT and anthracyclines. **a**, Schematic representation of experimental design for STING antagonist H-151 treatments following cardiac RT relative to study endpoints. **b**, Irf7 and Cxcl10 gene expression in cardiac fibroblasts isolated from *Sting*^*+/+*^ after cardiac RT or sham treatment and subsequent treatment with H-151 or vehicle, measured by qRT-PCR. **c**, Ejection fraction as measured by echocardiography in *Sting*^*+/+*^ mice 3 months after cardiac RT or shame treatment and subsequent treatment with either STING antagonist H-151 or vehicle until the time of echocardiography. **d**, Schematic representation of experimental design for STING antagonist H-151 treatments in mice treated with doxorubicin to induce cardiac toxicity. **e**, Ejection fraction of *Sting*^*+/+*^ mice 14 days after completing 5 weekly doses of doxorubicin or vehicle and concurrent H-151 of vehicle until the time of echocardiography. *p<0.05, **p<0.01, ***p<0.001, ****p<0.0001.

Our findings provide new mechanistic understanding of cardiac toxicity following DNA damaging cancer therapy. These results suggest that unrepaired DNA damage following radiation therapy or anthracyclines drive cytosolic mis-localization of DNA in fibroblasts weeks after DNA damage, which in turn is detected by cGAS and leads to the activation of the type I interferon response. The lag in between the DNA damage event and the induction of cGAS-STING dependent signaling is likely related to a requirement for fibroblasts to undergo cell cycling prior to cytosolic mis-localization of genomic DNA as previously reported ^24^. Ultimately, the chronic inflammation that results from this activation leads to cardiac mortality months after the initial insult. Further studies will need to determine whether these results in cardiac toxicity are part of a more generalized mechanism of delayed normal tissue injury following DNA damaging cancer therapies since targeting of the cGAS-STING pathway holds great potential as a treatment to prevent normal tissue toxicity and improve the therapeutic ratio of DNA damaging cancer treatments. Indeed, these data indicate that clinical trials utilizing cGAS-STING antagonists to prevent cardiac toxicity following DNA damaging cancer therapy are warranted. Finally, our results demonstrate the consequences of pathogenic cGAS-STING signaling in normal tissues which may have implications for drugs seeking to exploit cGAS-STING activation to treat cancer.

## Supporting information

Shamseddine et al Supplement

## Acknowledgments

We thank Simon Powell and Azusa Tanaka for careful reviews of the manuscript and helpful discussions. This work was supported by funding from Department of Defense Discovery Award W81XWH1910002 (SHP, SB, AMS), National Institutes of Health R35 GM 124909 (AMS), MSKCC Imaging and Radiation Sciences grant (SHP, SB, AMS), MSKCC Chanel Endowment for Survivorship Research Grant (AFY), and MSKCC Cancer Center Support Grant P30 CA 008748.

## Author contributions

Conceptualization: SHP and AMS

Methodology: AS, SHP, ATC, AFY, AP, NDS, SFB, AMS

Investigation: AS, SHP, VC, MA, MDB, JL, NDS, CTD, LVR, KCO, JEL, RMS, AFY, AMS

Visualization: AS, SHP, NDS

Funding acquisition: SHP, SFB, AFY, AMS

Project administration: AMS

Supervision: SFB and AMS

Writing – original draft: AS and AMS

Writing – review & editing: All authors

## Data and materials availability

Bulk RNA sequencing and single cell RNA sequencing data will be deposited in GEO. All other data are available in the main text or supplementary materials.

## Supplementary Materials

**Materials and Methods**

**Supplementary Figures 1-4**

**Supplementary Tables 1-2**

