## Supplementary material for "Innate immune signaling drives late cardiac toxicity following DNA damaging cancer therapies": Shamseddine et al Supplement

### Supplementary Materials

#### Materials and Methods

Mice, reagents and injections

Animal housing, handling and all procedures were approved by MSKCC ethical committees and were performed in accordance to IACUC guidelines. All mice strains were purchased from the Jackson laboratories, namely C57BL/6J (Cat#: 000664), and *Sting*$^{gt}$ (Cat#: 017537), *Cgas*$^{tm1d(EUCOMM)Hmgu}$ (Cat#: 026554, referred to as *Cgas*$^{-/-}$) and *Mavs*$^{tm1Zjc}$ (Cat#: 008634, referred to as *Mavs*$^{-/-}$).

Pharmaceutical grade doxorubicin was purchased from MSKCC pharmacy at a concentration of 2 mg/mL in PBS. For experiments mice were weighted prior to each injection and subsequently injected with 5 mg/kg doxorubicin or vehicle (PBS) weekly for 5 weeks to a total dose of 25 mg/kg. For inhibitor treatments, mice were injected intraperitoneally with daily 750 nmol of H151 (Tocris, Cat #:6675) in 200 uL PBS 0.1% Tween-80 or vehicle and 750 nmol of C176 STING (Tocris, Cat#: 7040) in 200 uL corn oil or vehicle starting at 7 days prior to radiation or doxorubicin treatment and continuing until experimental endpoint.

Radiation treatments

Mice aged 8-10 weeks were anesthetized using 3% isoflurane during induction and 2% during maintenance. X-Rad225Cx small animal irradiator (Precision X-ray) was used to administer either 12 or 20Gy of radiation to the heart using 225kVp X-rays and a 0.9mm half value layer Cu beam and a 10 mm circular collimator. The isocenter was placed at the center of the heart as determined by a cone beam CT. The 12 Gy treatment was delivered using parallel opposed obliques while the 20 Gy treatment was delivered using a 4 field box. Both treatments were delivered at a dose rate of 400 cGy/min. After analysis of dosimetry using a standard radiation protocol on multiple animals, total dose varied <5% across animals despite sex and age differences; therefore, beam angles and radiation plan were not adjusted for sex or weight. Sham irradiation involved all steps including sedation, imaging, and recovery, except actual radiation delivery.

Survival analyses

Mice were euthanized when showing evidence of overt heart failure including hunching, rapid breathing, lethargy and abdominal distention. Kaplan Meier curves were plotted using Graphpad prism. Hearts were paraffin embedded and examined by pathologist to confirm histopathologic evidence of radiation induced cardiac toxicity.

Heart cell separation

Mice were euthanized at experimental timepoints per IACUC guidelines. To reduce variability and red blood cell contamination as well as RNA degradation, upon entering the thoracic cavity, cold PBS was perfused immediately before further cardiac extraction. The great vessels were cut at their interface to the heart, and pericardium and surrounding adipose were discarded.

Endothelia and fibroblasts

Hearts were harvested and digested with gentleMACS C Tube (Miltenyi, Cat#: 130-093-237) and the neonatal heart dissociation kit (Miltenyi, Cat#: 130-098-373) in a gentleMACS Octo Dissociator with heaters according to manufacturer's protocol. Following dissociation, debris removal and red blood cell lysis was performed using debris removal solution (Miltenyi, Cat#: 130-109-398) and red blood cell lysis solution (Miltenyi, Cat#: 130-094-183) respectively according to manufacturer's protocol. Lysates were subsequently sequentially immunoprecipitated on an OctoMACS magnetic separator in MS columns (Miltenyi, Cat#: 130-042-201) using mouse CD45 (Miltenyi, Cat#: 130-052-301) to deplete leukocytes, CD31 (Miltenyi, Cat#: 130-097-418) and CD90.2 (Miltenyi, Cat#: 130-121-278) microbeads according to manufacturer's protocol.

Myocytes

Following acute heart extraction, myocytes were isolated utilizing an enzymatic isolation protocol via manufacturer instructions (Adumyt Non-perfusion Adult Cardiomyocyte Isolation, Cellutron Life Technologies). Because cells were not intended for culture, a Langendorf perfusion system was not utilized.

RNA isolation, purification and cDNA generation

RNA isolation was performed using RNeasy Mini Kit (Qiagen, Cat#: 74106) according to manufacturer's protocol. Following RNA isolation, RNA was purified using RNAClean XP (Beckman Coulter, Cat#: A63987) per manufacturer's instruction. Genomic DNA was digested using ezDNAse Enzyme (ThermoFisher, Cat#: 11756050) for 2 minutes at 37 degrees and cDNA was generated using SuperScript IV Vilo Master Mix (ThermoFisher, Cat#: 11756050) according to manufacturer's protocol.

Quantitative Reverse Transcriptase PCR (qRT-PCR)

RT-qPCR was performed using TaqMan$^{TM}$ Fast Advanced Master Mix (ThermoFisher, Cat#: 4444556) on an Applied Biosytems QuantStudio 6 Flex Real-Time PCR system using the indicated primers: Irf7 (ThermoFisher, Cat#: Mm00516793_g1), Cxcl10 (ThermoFisher, Cat#: Mm00445235_m1), GAPDH (ThermoFisher, Cat#: Mm99999915_g1). Expression was normalized to GAPDH (mice).

RNA sequencing and analysis

RNA sequencing on poly-A selected RNA and was performed by Novogene on the Illumina NovaSeq 6000 Sequencing System. Samples were quantitated for total RNA and subsequent steps were carried out by Novogene, including library preparation and paired-end 150 bp sequencing. Bioinformatics analyses including quality control and DEseq were performed by Basepair with FDR corrections for multiple tests. FDR < 0.05 was considered a statistically significant difference. Additional analysis for GO enrichment was performed using the web based software Panther (geneontology.org).

Single-cell transcriptome sequencing

Dissociated, sorted cells were stained with Trypan blue and the Countess II Automated Cell Counter (ThermoFisher) was used to assess both cell number and viability (74%-82%). Following QC, the single cell suspension was loaded onto Chromium Chip B (10X

Genomics PN 2000060) and GEM generation, cDNA synthesis, cDNA amplification, and library preparation of 4,500-9,300 cells proceeded using the Chromium Single Cell 3' Reagent Kit v3 (10X Genomics PN 1000075) according to the manufacturer's protocol. cDNA amplification included 11 cycles and 44-87ng of the material was used to prepare sequencing libraries with 12 cycles of PCR. Indexed libraries were pooled equimolar and sequenced on a NovaSeq 6000 in a PE28/91 paired end run using the NovaSeq 6000 SP, S1, or S2 Reagent Kit (100 cycles) (Illumina). An average of 201 million paired reads was generated per sample

Single cell transcriptome analysis

The raw sequence data (FASTQ files) were first processed with 10X Genomics CellRanger (ver 5) software to compute the cell/barcode by gene count matrices. The command used was: cellranger count and the reference database was refdata-gex-mm10-2020-A. The count matrix was then processed with a series of R scripts (R version 3.6.1) using the Seurat (version 3.2.2) R library. The workflow used was included standard processing with the SCTransform method for normalization and sample integration. First after reading in the 10X data the cells in each sample were filtered to remove those that had any of the following qc measures: less than 1,500 detected genes per cell or less than 5,000 UMI's per cell or greater than 10% of the cells reads mapping to mitochondrial genes. Cells that failed any of these qc tests were removed from the analysis. Next each cell was scored for its cell cycle phase using Seurat's CellCycleScoring function. Note the Seurat library only has cell cycle genes for human so we obtained a mouse version using R's biomaRt packaged to convert between human and mouse homologues.

At this stage all samples were processed independently. The next step was to integrate all of them into one unified dataset using Seurat's integration workflow; with SCTransform to normalize the data before integration. This first step both normalizes and regresses out the cell cycle component. We then followed the rest of the integration workflow from the Seurat vignette.

At this point we made two passes through the following pipeline. In the first pass the data was clustered and cell types identified using custom gene lists and Seurat's AddModuleScore function. Clusters that contained irrelevant biological cell types were then filtered out and we repeated the entire processing pipeline with the reduced set of cells. The cell type filtering was down after the integration step. We ran the PCA projection (RunPCA), computed the UMAP embedding (RunUMAP) using the first 20 PCA components. The data was then clustered using the FindNeighbors/FindClusters functions with a resolution of 0.2 in the clustering step.

We then found cluster specific marker genes and re-computed the cell type identifications as described above.

To compute differential gene expression, we use the Muscat package (ver 1.5.0, for muscat ver 4.0 of R was used for compatibility reasons). Specifically, we used the pseudo-bulk method with edgeR.

Cardiac 2D echocardiography

Mice were anesthetized using 3% isoflurane during induction and 2% during maintenance. Mice were placed on a heated stage to 37C and 2D echocardiograms were obtained using FujiFilm-Vevo 2100 micro-Ultrasound in B mode. Parasternal long axis views were utilized for ejection fraction and fractional shortening analysis using the Vivo lab LV trace tool.

Sample preparation for Immunofluorescence

The immunofluorescence detection of cGAS was performed at Molecular Cytology Core Facility of Memorial Sloan Kettering Cancer Center, using Discovery XT processor (Ventana Medical Systems). After 32 minutes of heat and CC1 (Cell Conditioning 1, Ventana cat#: 950-500) retrieval, the tissue sections were blocked first for 30 mins in Background Blocking reagent (Innovent, catalog #: NB306). A rabbit monoclonal antibody to cGAS (Cell Signaling, cat#: 31659) was used at a concentration of 0.5 ug/mL. The incubation with the primary antibody was done for 5 hours, followed by 60 minutes incubation with biotinylated goat anti-rabbit IgG (Vector labs, cat#: PK6101) at a concentration of 5.75 mg/mL., followed by Blocker D, Streptavidin-HRP and TSA Alexa 488 (Life Tech, cat#: B40932) for 16 minutes.

The next day, a rabbit monoclonal Vimentin antibody (Cell Signaling Cat#: 5741) was used in a 0.045 ug/mL concentration. The incubation with the primary antibody was done for 5 hours, followed by a 60 minutes incubation with a goat anti-rabbit IgG (Vector labs, cat#: PK6101) to a 5.75 mg/mL concentration. Blocker D, Streptavidin-HRP and Tyramide-CF594 (Biotium, Cat#: 92174) were prepared according to manufacturer instructions in a 1:2000 dilution and incubated for 16 minutes.

All slides were counterstained with 5 ug/mL DAPI (dihydrochloride(2-(4-Amidinophenyl)-6-indolecarbamidine dihydrochloride) (Sigma, Cat#: D9542) for 5 minutes at room temperature, mounted with anti-fade mounting medium Mowiol (Mowiol 4-88 CalbioChem code: 475904) and coverslip was added.

Immunofluorescence image acquisition

The images were acquired with an inverted Zeiss LSM880 (Carl Zeiss Microscopy GmbH, Carl-Zeiss-Straße 22, 73447 Oberkochen, Germany) equipped with an Airy Scan detector (gain 850, digital gain 1) in super resolution mode. The objectives used for acquisition were a 20x air immersion (Plan-Apochromat 20x/0.8 M27) and a 63x oil immersion (Plan-Apochromat 63x/1.4 Oil DIC M27). With the 63x magnification we used a pixel size of 32 nm and dwell time of 19.8 us for the zoomed images and a pixel size and dwell time of 106 nm and 39.6 us, respectively, for the bigger field of view. With the 20x objective we used a pixel dwell time of 39.6 us and a pixel size of 334 nm. We used the same optical configuration, splitting the three laser lines (405nm at 0.7% power, 488nm at 0.4% and

561nm at 0.5%) with a 488/561/633 MBS and an invisible light beam splitter at 405. In detection we applied a band pass filter 495-550 nm and a long pass 570 nm.

For quantification, we acquired 10 images for each sample with the same optical configuration and 63x objective, but with pixel size of 265 nm and pixel dwell time of 1.24 us. Z-stacks were acquired when the thickness of the sample required it, with a spacing of 500 nm between the stacks, and a maximum intensity projection was performed before quantification using the Zeiss ZEN black software.

All the images were processed using the feature AiryScan processing in the Zeiss ZEN software for Image Acquisition and Analysis, using default parameters. Further image processing was performed in ImageJ (Rasband, W.S., ImageJ, U.S. National Institutes of Health, Bethesda, Maryland, USA, https://imagej.nih.gov).

MSKCC IRB 14-099 Clinical Trial and Biospecimens

A prospective cohort of 80 women with non-metastatic breast cancer patients who were planned for treatment with anthracycline-based polychemotherapy followed by trastuzumab were enrolled. All patients underwent routine 2D and speckle tracking echocardiography at baseline (pre-anthracycline) and, after the completion of anthracyclines (2months) and thereafter at 3 months intervals until one year. Blood was collected at the pre-anthracycline and the 2 months timepoint and analyzed for biomarkers.

Cytokine Analyses

Mice were euthanized at indicated timepoint and intracardiac puncture was performed using a 21 Gauge needle. Blood was collected at room temperature and held upright before for 30 minutes. Blood was spun at 1500 g for 15 mins and serum was collected by collecting supernatant. For ELISA, 100 uL of serum was run in duplicate using the mouse CXCL10 sandwich ELISA (R&D systems, Cat# DY466) according to manufacturer's protocol. For Luminex, 25 uL of serum was run in duplicate using the CXCL10 mouse Procarta simplex kit (ThermoFisher Cat# EPX01A-26018-901) according to manufacturer's protocol. Samples were red on a Flexmap 3D machine.

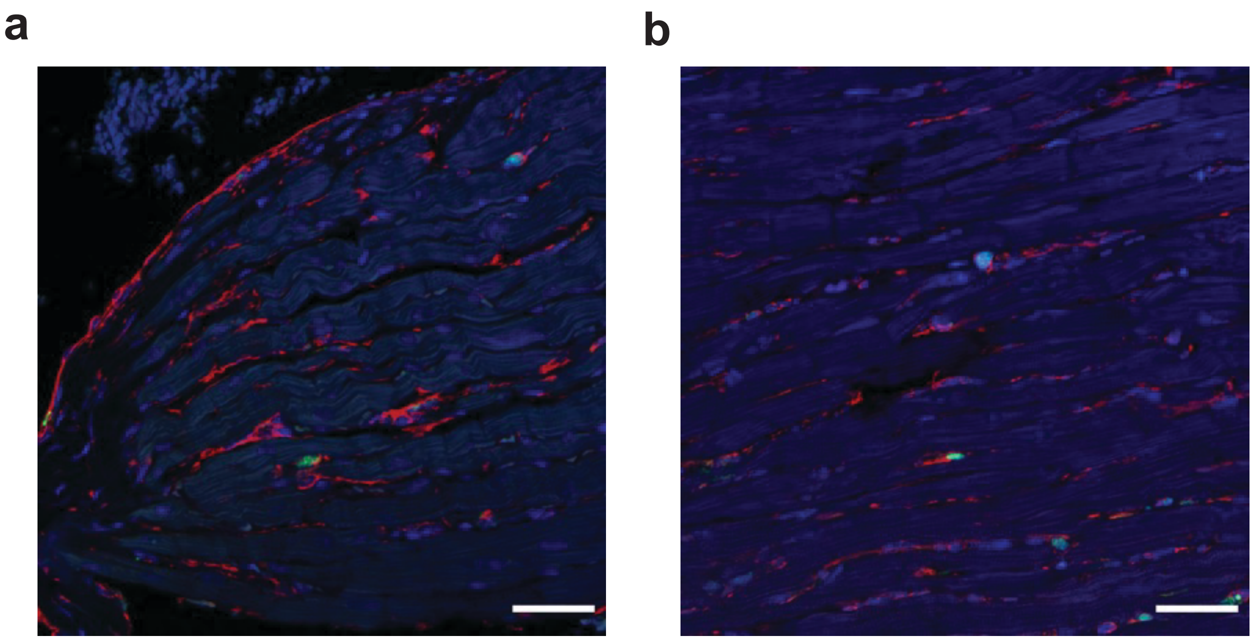


**Supplementary Figure 1.** Low Magnification (40X) immunofluorescence images of hearts sections 28 days after sham treatment (**a**) and cardiac RT (**b**). (cGAS - Green, Vimentin - Red, DAPI - Blue). Related to **Figure 2**.

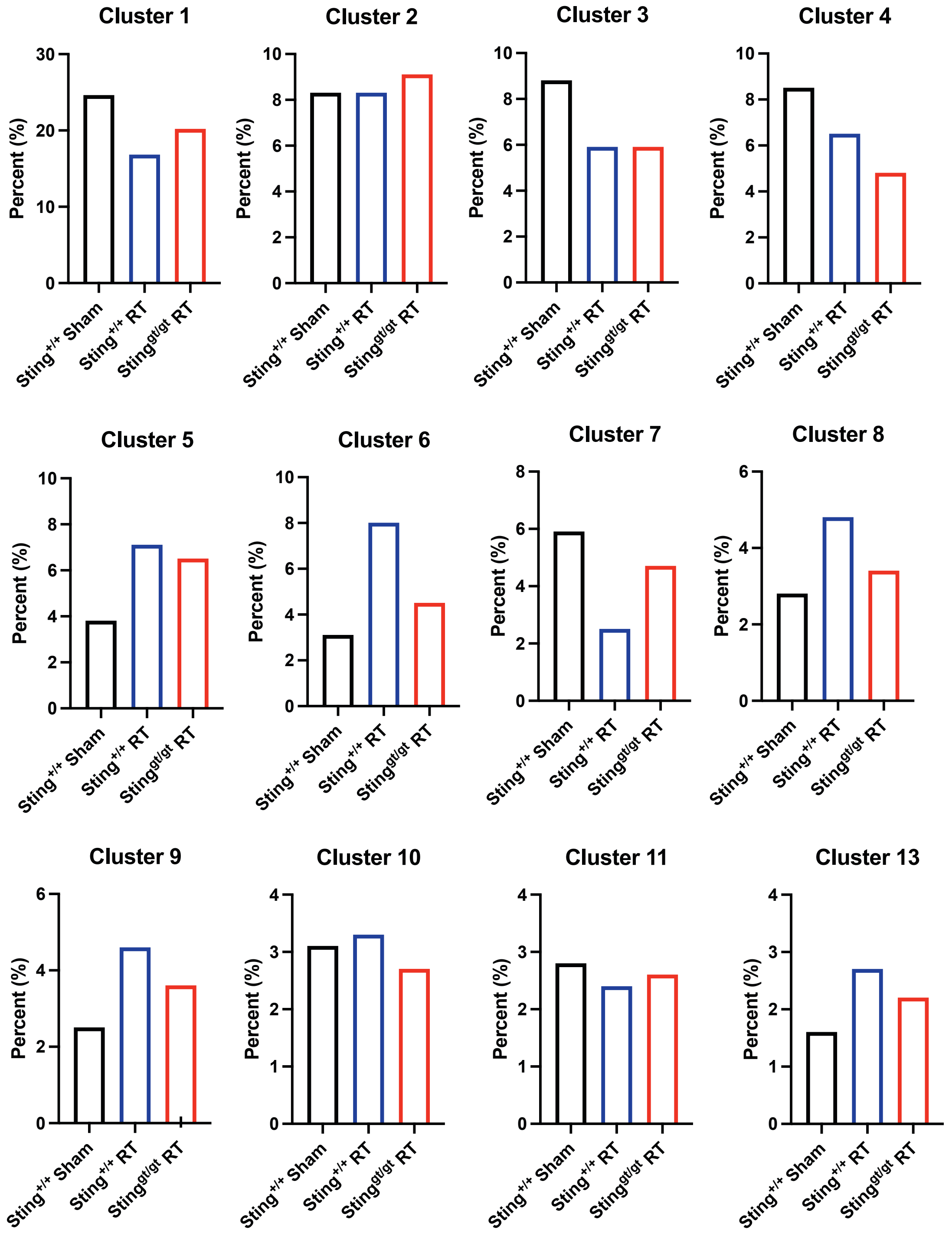


**Supplementary Figure 2.** Each cluster as identified in UMAP as a percentage of the total cell population in *Sting*$^{+/+}$ after 28 days after sham treatment, *Sting*$^{+/+}$ mice 28 days after cardiac RT, and *Sting*$^{gt/gt}$ mice 28 days after cardiac RT. Related to **Figure 3**.

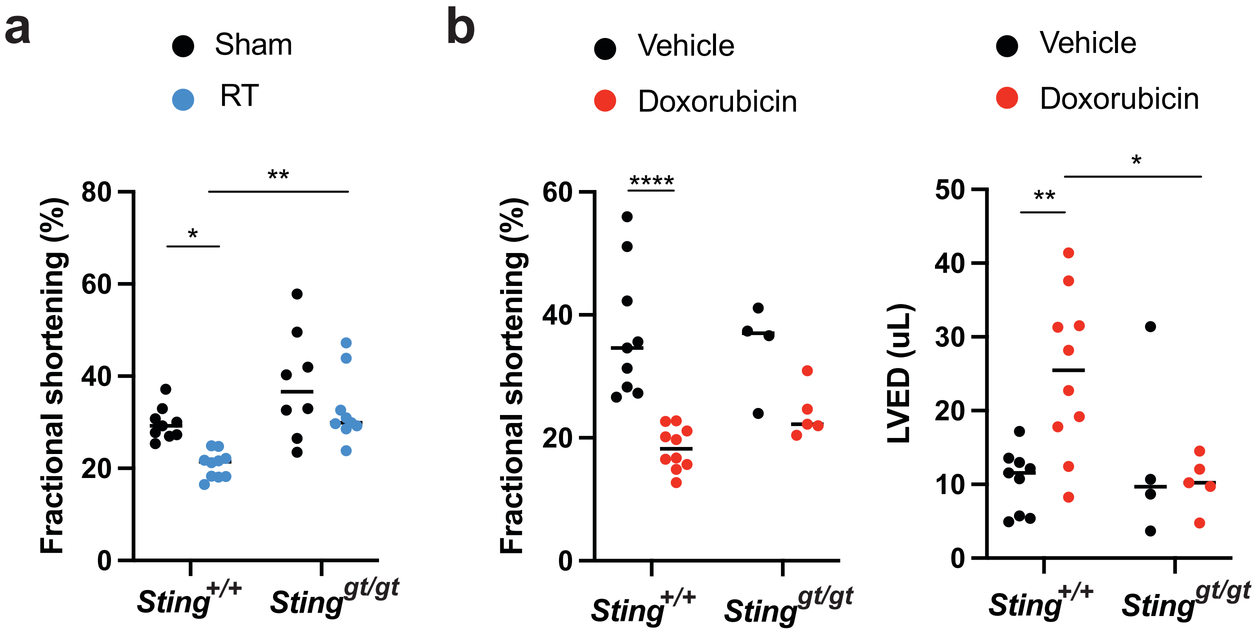


**Supplementary Figure 3. a,** Fractional shortening in sham and RT-treated mice of indicated genotypes 3 months after cardiac RT or sham treatment. **b,** Fractional shortening and Left ventricular end diastolic volume in mice of indicated genotypes 14 days after completing 5 weekly doses of doxorubicin or vehicle. (*p<0.05, **p<0.01, ***p<0.001, ****p<0.0001). Related to Fig. 3.

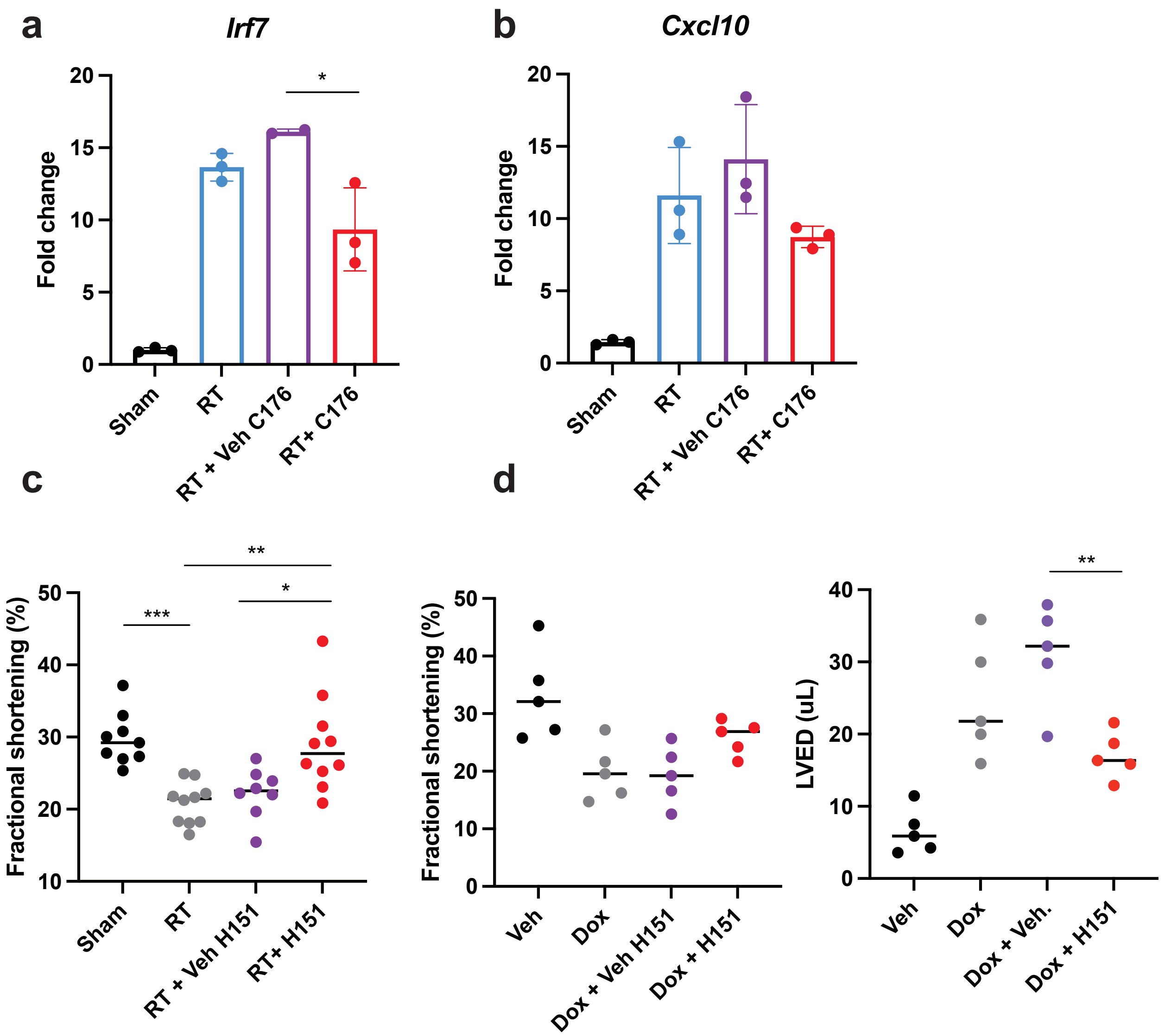


**Supplementary Figure 4. a-b**, qRT-PCR for expression of *Irf7* (**a**) and *Cxcl10* (**b**) in cardiac fibroblasts 28 days after cardiac RT or sham and 7 days of treatment with STING antagonist C-176 or vehicle. **c,** Fractional shortening as measured by echocardiography in *Sting*$^{+/+}$ mice 3 months after cardiac RT or sham treatment and subsequent treatment with either STING antagonist H-151 or vehicle until the time of echocardiography. (**d-e**) Fractional shortening (**d**) and LVED (**e**) of *Sting*$^{+/+}$ mice 14 days after completing 5 weekly doses of doxorubicin or vehicle and concurrent H-151 of vehicle until the time of echocardiography.

**Supplementary Table 1.** Specific GO terms corresponding to rows in Figure 1c.

| | **GO TERM** |
|---|---|
| **1** | Positive regulation of cysteine-type endopeptidase activity involved in apoptotic signaling |
| **2** | Release of cytochrome c from mitochondria |
| **3** | Positive regulation of release of cytochrome c from mitochondria |
| **4** | Intrinsic apoptotic signaling pathway in response to endoplasmic reticulum stress |
| **5** | Leukocyte apoptotic process |
| **6** | DNA damage response, signal transduction by p53 class mediator resulting in cell cycle arrest |
| **7** | Intrinsic apoptotic signaling pathway in response to DNA damage by p53 class mediator |
| **8** | Embryonic viscerocranium morphogenesis |
| **9** | Adhesion to symbiont host cell |
| **10** | Defense response to protozoan |
| **11** | T cell homeostasis |
| **12** | T-helper cell differentiation |
| **13** | Mitotic spindle midzone assembly |
| **14** | Regulation of attachment of spindle microtubules to kinetochore |
| **15** | Positive regulation of fibroblast proliferation |
| **16** | Regulation of microtubule polymerization or depolymerization |
| **17** | Positive regulation of type I interferon-mediated signaling pathway |
| **18** | Negative regulation of type I interferon-mediated signaling pathway |
| **19** | Type I interferon signaling pathway |
| **20** | Positive regulation of interferon-beta production |
| **21** | Positive regulation of IRE1-mediated unfolded protein response |
| **22** | Negative regulation of endoplasmic reticulum calcium ion concentration |

**Supplementary Table 2.** Genes Most Significantly associated with each scRNA-seq cluster in Figure 3a and Supplementary Figure 2.

| 0 | 1 | 2 | 3 | 4 | 5 | 6 | 7 | 8 | 9 | 10 | 11 | 12 | 13 |
|---|---|---|---|---|---|---|---|---|---|---|---|---|---|
| H2-Ab1 | Pf4 | Plac8 | Hdc | Trbc2 | Fn1 | Cd79a | Il1b | Cd209a | H2afz | Cd79a | Ctsd | Ifit3 | Xcr1 |
| H2-Eb1 | Timp2 | Ifitm3 | Il1r2 | Ms4a4b | Ear2 | Ly6d | Csf3r | Ifi30 | Birc5 | Ebf1 | Ctsb | Ifit2 | Clec9a |
| H2-Aa | Selenop | Tmsb10 | Il1b | Cd3e | Tnip3 | Cd79b | Msrb1 | Wnt11 | Ccna2 | Igkc | Cd63 | Isg15 | Sept3 |
| Cd74 | Folr2 | Chil3 | Cebpb | Cd3d | Retnla | Ms4a1 | S100a8 | Ddr1 | Nusap1 | Ly6d | Prdx1 | Ifit3b | Cd24a |
| H2-DMb1 | Mrc1 | Msrb1 | S100a11 | Nkg7 | Mmp12 | Igkc | S100a9 | Bhlhe40 | Mki67 | Ms4a1 | Cstb | Fcgr1 | Tmsb10 |
| Cxcl16 | Stab1 | Napsa | S100a9 | Cd3g | Furin | Iglc2 | Hdc | Klrd1 | Pclaf | Ighd | Lgals3 | Oasl2 | Itgae |
| | Cbr2 | Ly6c2 | S100a8 | Il2rb | Lpl | Ebf1 | Gsr | Klrb1b | Cdca3 | Fcmr | Lipa | Bst2 | Wdfy4 |
| | Gas6 | Ifitm6 | Csf3r | Trac | Lyz1 | Mzb1 | S100a11 | Tnip3 | Prc1 | | Creg1 | Ifit1 | Ifi205 |
| | Lyve1 | S100a4 | Msrb1 | Trbc1 | | Ighm | Cytip | Traf1 | Cdk1 | | Lpl | Irf7 | Naaa |
| | Apoe | Hp | G0s2 | Gzmb | | H2-DMb2 | Il1r2 | Etv3 | Stmn1 | | Igf1 | Ifi204 | Ppt1 |
| | F13a1 | Apoc2 | Trem1 | Txk | | Iglc3 | G0s2 | | Cdca8 | | Gdf15 | Cmpk2 | Irf8 |
| | Fcgrt | Gsr | Retnlg | Ets1 | | Pou2af1 | Mxd1 | | Top2a | | Trem2 | Usp18 | Gm2a |
| | Ltc4s | Ms4a4c | Mxd1 | Ptprcap | | Cd19 | Hp | | Tpx2 | | Hmox1 | Phf11b | Jaml |
| | Rnase4 | Smpdl3a | Clec4d | Gimap6 | | Ptprcap | Lmnb1 | | Cenpf | | Gclm | Rsad2 | Naga |
| | Serpinb6a | Fn1 | Slpi | Gimap4 | | Fcmr | Cd44 | | Ube2c | | Slc40a1 | Ifi47 | Cst3 |
| | Fcna | Prdx5 | Acod1 | Il7r | | Ly6a | Trem1 | | Cks1b | | Clec4n | Rtp4 | Ckb |
| | Ccl24 | Mgst1 | Cxcr2 | Gzma | | Scd1 | Cebpb | | Ccnb2 | | Cd36 | Slfn5 | Plbd1 |
| | Cfh | Gngt2 | Slc7a11 | Satb1 | | Fcrla | Slpi | | Tubb5 | | Mmp12 | Phf11d | Trim35 |
| | Ninj1 | Thbs1 | Lmnb1 | Ccl5 | | Cd37 | Clec4d | | Racgap1 | | Spp1 | Iigp1 | |
| | Wfdc17 | | Mmp9 | Vps37b | | Pax5 | Btg1 | | Smc2 | | Fabp5 | Ifi211 | |
| | Ccl8 | | Hp | AW112010 | | Iglc1 | Samsn1 | | Tuba1b | | Ftl1 | Ifi203 | |
| | Cd163 | | Wfdc21 | | | Bank1 | Slc7a11 | | Hist1h2ae | | Fth1 | Parp14 | |
| | | | Srgn | | | Ccr7 | Retnlg | | Kif23 | | | Ccl12 | |
| | | | Slc16a3 | | | H2-Ob | Tnfaip2 | | Cdkn2c | | | Pnp | |
| | | | Lrg1 | | | Cd24a | Acod1 | | H2afv | | | Cxcl10 | |
| | | | Pglyrp1 | | | Cd2 | Grina | | Tmpo | | | Ms4a4c | |
| | | | Grina | | | Ets1 | Cxcr2 | | Hist1h2ap | | | | |
| | | | Ifitm1 | | | Ighd | Slc16a3 | | Smc4 | | | | |
| | | | Tnfaip2 | | | | Pglyrp1 | | Hmgb2 | | | | |
| | | | Lcn2 | | | | Pim1 | | Hmgn2 | | | | |
| | | | Ccr1 | | | | Plaur | | H2afx | | | | |
| | | | Rdh12 | | | | Itgal | | Hmgb1 | | | | |
| | | | Clec4e | | | | Mmp9 | | Nucks1 | | | | |
| | | | Plaur | | | | Mcl1 | | Arl6ip1 | | | | |
| | | | Ptgs2 | | | | Ets2 | | | | | | |
| | | | Thbs1 | | | | Sorl1 | | | | | | |
| | | | Cstdc4 | | | | Lrg1 | | | | | | |
| | | | | | | | Taldo1 | | | | | | |
| | | | | | | | Gcnt2 | | | | | | |
| | | | | | | | Ifitm1 | | | | | | |
| | | | | | | | Ccr1 | | | | | | |
| | | | | | | | Ndel1 | | | | | | |
| | | | | | | | S100a6 | | | | | | |
| | | | | | | | Cxcr4 | | | | | | |
| | | | | | | | Mcemp1 | | | | | | |
| | | | | | | | Arg2 | | | | | | |
| | | | | | | | Clec4e | | | | | | |
| | | | | | | | Lcn2 | | | | | | |
| | | | | | | | Wfdc21 | | | | | | |
| | | | | | | | Stk17b | | | | | | |
| | | | | | | | Srgn | | | | | | |
| | | | | | | | Ptgs2 | | | | | | |
| | | | | | | | Cd300lf | | | | | | |
| | | | | | | | Litaf | | | | | | |
| | | | | | | | Fosl2 | | | | | | |
| | | | | | | | Trib1 | | | | | | |
| | | | | | | | Sgms2 | | | | | | |
| | | | | | | | Tgm2 | | | | | | |
| | | | | | | | Gda | | | | | | |
| | | | | | | | Stfa2l1 | | | | | | |
| | | | | | | | Tpd52 | | | | | | |
| | | | | | | | Rdh12 | | | | | | |
| | | | | | | | Cxcl2 | | | | | | |
| | | | | | | | Thbs1 | | | | | | |
| | | | | | | | Asprv1 | | | | | | |
| | | | | | | | Cstdc4 | | | | | | |
| | | | | | | | Hcar2 | | | | | | |
| | | | | | | | Slc2a3 | | | | | | |
| | | | | | | | Ankrd33b | | | | | | |
| | | | | | | | Pygl | | | | | | |
| | | | | | | | Mmp8 | | | | | | |
| | | | | | | | Adam8 | | | | | | |
| | | | | | | | Plk3 | | | | | | |
